# Calmodulin enhances mTORC1 signaling by preventing TSC2-Rheb binding

**DOI:** 10.1101/2024.08.19.608556

**Authors:** Yuna Amemiya, Yuichiro Ioi, Masatoshi Maki, Hideki Shibata, Terunao Takahara

## Abstract

The mechanistic target of rapamycin complex 1 (mTORC1) functions as a master regulator of cell growth and proliferation. We previously demonstrated that intracellular calcium ion (Ca^2+^) concentration modulates the mTORC1 pathway via binding of the Ca^2+^ sensor protein calmodulin (CaM) to tuberous sclerosis complex 2 (TSC2), a critical negative regulator of mTORC1. However, the precise molecular mechanism by which Ca^2+^/CaM modulates mTORC1 activity remains unclear. Here, we performed a binding assay based on nano-luciferase reconstitution, a method for detecting weak interactions between TSC2 and its target, Ras homolog enriched in brain (Rheb), an activator of mTORC1. CaM inhibited the binding of TSC2 to Rheb in a Ca^2+^-dependent manner. Live-cell imaging analysis indicated increased interaction between the CaM-binding region of TSC2 and CaM in response to elevated intracellular Ca^2+^ levels. Furthermore, treatment with carbachol, an acetylcholine analog, elevated intracellular Ca^2+^ levels, and activated mTORC1. Notably, carbachol-induced activation of mTORC1 was inhibited by CaM inhibitors, corroborating the role of Ca^2+^/CaM in promoting the mTORC1 pathway. Consistent with the effect of Ca^2+^/CaM on the TSC2-Rheb interaction, increased intracellular Ca^2+^ concentration promoted the dissociation of TSC2 from lysosomes without affecting Akt-dependent phosphorylation of TSC2, suggesting that the regulatory mechanism of TSC2 by Ca^2+^/CaM is distinct from the previously established action mechanism of TSC2. Collectively, our findings offer mechanistic insights into TSC2–Rheb regulation mediated by Ca^2+^/CaM, which links Ca^2+^ signaling to mTORC1 activation.

## Introduction

The mechanistic target of rapamycin complex 1 (mTORC1) is a vital regulator of cell growth and proliferation, controlling both anabolic and catabolic processes. This kinase complex comprises three essential components: mTOR kinase, Raptor, and mLST8, and it integrates extracellular and intracellular signals, including amino acids, insulin, and various stresses, to regulate cell growth (1–3). mTORC1 activity is regulated by two distinct small GTPases: Ras-related GTP binding (Rag) GTPase and Ras homolog enriched in brain (Rheb) GTPase. Rag GTPases, which form heterodimers of RagA or RagB with RagC or RagD, reside on the cytosolic surface of lysosomes through the scaffold protein complex Ragulator. The nucleotide state of Rag GTPases is primarily regulated by amino acid availability (4), and active Rag GTPases recruit mTORC1 to lysosomes by binding to the Raptor (5, 6). Thus, lysosomes are considered major cellular sites of mTORC1 activation.

On the other hand, Rheb GTPase is believed to bind to mTORC1, inducing an active conformational change and acting as a direct activator of mTORC1 (7). While the exact cellular localization of Rheb remains unclear (8–10), some studies suggest its lysosomal localization potentially facilitates mTORC1 activation (11, 12). Growth factors, such as insulin, activate Rheb, leading to mTORC1 activation (11). In addition, recent studies have shown that amino acids also play a crucial role in regulating Rheb to modulate mTORC1 signaling (13–16). The nucleotide state of Rheb is primarily regulated by the tuberous sclerosis complex (TSC), which acts as a GTPase-activating protein (GAP). The TSC complex, composed of TSC1, TSC2, and Tre2-Bub2-Cdc16-1 domain family member 7 (TBC1D7), is a potent negative regulator of mTORC1 signaling (1). Within this complex, TSC2 functions as a Rheb-GAP while other subunits stabilize it (17–22). Mutations in *TSC1* and *TSC2* cause the condition known as tuberous sclerosis complex, which is characterized by systemic benign tumors and epilepsy (23). Upon insulin stimulation, Akt is activated through phosphatidylinositol-3-kinase (PI3K)–mTOR complex 2 (mTORC2) signaling, which inactivates TSC2 and leads to Rheb GTPase activation. Akt directly phosphorylates TSC2 (24). Although several potential roles of TSC2 phosphorylation by Akt have been proposed (11), the most persuasive role is in regulating the cellular localization of TSC2, particularly its lysosomal localization and dynamics (11, 15, 25, 26). Moreover, recent findings indicate that TSC2 localization is affected by stress, including amino acid deprivation and hyperosmotic stress (27). However, the mechanisms by which TSC2 regulates Rheb beyond its phosphorylation-dependent subcellular localization remain unclear.

Calcium ions (Ca^2+^) act as a crucial second messenger in various biological processes. mTORC1 signaling is also modulated by intracellular Ca^2+^ mobilization (28). A link between intracellular Ca^2+^ levels and mTORC1 signaling in physiological and pathological cardiac hypertrophy has also emerged recently (28, 29). Previously, we and others reported that the addition of amino acids causes an increase in intracellular Ca^2+^, which is sensed by CaM, leading to mTORC1 activation in amino acid-starved cells (16, 30, 31). Ca^2+^/CaM binds to human vacuolar protein sorting 34, activating mTORC1 (30). As another mechanism of the Ca^2+^/CaM effect on mTORC1 signaling, we demonstrated that Ca^2+^/CaM binds to TSC2, leading to mTORC1 activation (16). However, the detailed molecular mechanism of this activation remains largely unclear.

In this study, we have investigated the mechanisms by which Ca^2+^/CaM binding to TSC2 modulates the association between TSC2 and Rheb. We have showed CaM can bind to the GAP domain of TSC2 in a Ca^2+^-dependent manner, which in turn induces disruption of the TSC2-Rheb association. We further found that the treatment of cells with carbachol, a Ca^2+^ mobilization agent, also enhances mTORC1 activity in a CaM-dependent manner, suggesting that Ca^2+^/CaM generally plays a key role in mTORC1 activation in response to the change in intracellular Ca^2+^ levels. Thus, our results revealed that Ca^2+^/CaM-mediated prevention of TSC2 binding to Rheb is a key mechanism linking the Ca^2+^ mobilization with subsequent mTORC1 activation.

## Results

### CaM disrupts TSC2–Rheb interaction in a Ca^2+^-dependent manner

We previously identified a CaM-binding region within the TSC2 GAP domain (16), which was consistent with earlier findings (32) (Fig. 1A). Notably, this CaM-binding region (residues 1717–1732) nearly coincides with one α-helix (residues 1716–1731) that forms a pair of α-helices proposed to facilitate the interaction between TSC2 and Rheb (21) (Fig. 1A). This observation led us to hypothesize that CaM binding to the α-helix of TSC2 may influence the TSC2–Rheb interaction, thereby leading to activation of mTORC1.

**Figure 1.**
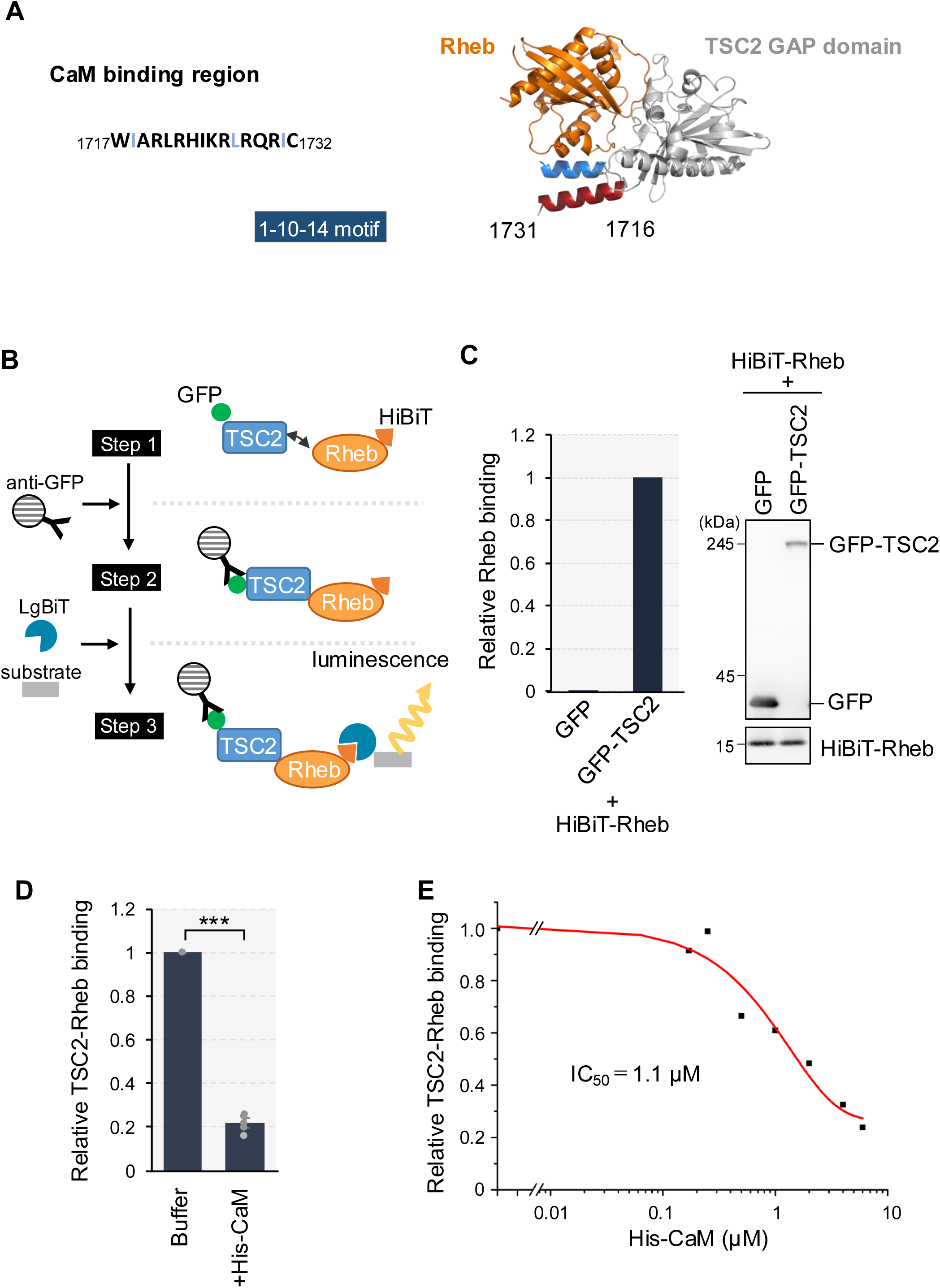
Ca^2+^ /CaM prevents TSC2-Rheb interaction. (A) The amino acids sequence for the CaM-binding region in TSC2 was shown, which matches the motif for the 1-10-14 type (49) (left). The predicted structure of Rheb and the GAP domain of TSC2 using AlphaFold2 is shown (right). The parallel α-helices which are thought to support the binding of Rheb with TSC2 are shown in blue and red (residues 1716-1731). α-helix (red) corresponds to the CaM-binding region. (B) Schematic presentation of the HiBiT lytic assay. The assay was performed in three steps. Strep-GFP-tagged TSC2 (GFP-TSC2) and HiBiT-tagged Rheb (HiBiT-Rheb) were mixed in vitro (Step 1). HiBiT-Rheb bound to GFP-TSC2 were purified by immunoprecipitation with anti-GFP antibody (Step 2). NanoLuc luciferase activity can be recovered by reconstitution of HiBiT with LgBiT, and then generates luminescence using the substrate (Step 3). The amounts of Rheb in GFP-TSC2 immunoprecipitates were detected as the luminescence intensity. (C) HEK293T cells were transiently transfected with the plasmid encoding Strep-GFP (GFP), Strep-GFP-TSC2 (GFP-TSC2) or HiBiT-Rheb. After 24 h of transfection, cells were lysed and respective lysates were mixed. Then, the cell lysates were subjected to immunoprecipitation using the anti-GFP antibody overnight, followed by the addition of Protein G magnetic beads. After washing the beads, the amounts of HiBiT and Strep were monitored as described in Experimental Procedures. The ratio of HiBiT-Rheb to GFP or GFP-TSC2 was shown (left). The samples were also analyzed with western blotting using the anti-GFP antibody and LgBiT blotting for detecting HiBiT-Rheb (right). (D) HEK293T cells were transiently transfected with the plasmid encoding Strep-GFP-TSC2 or HiBiT-Rheb. After 24 h of transfection, cells were lysed and respective cell lysates were mixed in the absence (Buffer) or presence of purified His-CaM (+His-CaM, 5.5 μM) together with 100 μM of CaCl_2_. Subsequent procedures were done as in (C). Graphs represent mean ± SEM of five independent experiments, One-way ANOVA with Tukey’s test,***p <0.001. (E) Assays were done as described in (D) using different amounts of His-CaM (0, 0.17, 0.25, 0.50, 1.0, 2.0, 4.0, 6.0 μM). Average results from two independent experiments were used to calculate the IC_50_ value.

To confirm this hypothesis, we assessed the binding between TSC2 and Rheb. To address the challenge of detecting weak TSC2-Rheb interactions using conventional immunoprecipitation (IP) followed by western blotting, we employed the HiBiT lytic assay, which is based on NanoLuc luciferase reconstitution and enables quantitative measurement of protein interactions, like our previous report (33). Strep-GFP-tagged TSC2 (GFP-TSC2) and HiBiT-tagged Rheb (HiBiT-Rheb) were separately expressed in HEK293T cells, respective cell lysates were mixed, and then subjected to immunoprecipitation (IP) using an anti-GFP antibody (Fig. 1B). Subsequently, the amount of HiBiT-Rheb within the purified immunoprecipitates was measured by adding the NanoLuc large fragment (LgBiT) to reconstitute NanoLuc by inducing the assembly of HiBiT with LgBiT. Specific binding between TSC2 and Rheb was successfully detected using the HiBiT lytic assay without degrading either protein (Fig. 1C).

To investigate the effect of CaM on the TSC2-Rheb interaction, purified recombinant His-tagged CaM (His-CaM) was added during this experiment. The addition of His-CaM decreased TSC2-Rheb interaction in the presence of 100 μM Ca^2+^ (Fig. 1D), indicating that CaM affects TSC2-Rheb association. We then assessed the inhibitory effect of Ca^2+^/CaM by adding varying amounts of His-CaM (Fig. 1E). The relative amount of Rheb bound to TSC2 was plotted as a function of His-CaM concentration, resulting in a dose-response inhibition curve. The calculated IC_50_ value of CaM for the TSC2-Rheb interaction was approximately 1.1 µM.

To determine whether Ca^2+^ is required for the effect of CaM on the TSC2-Rheb interaction, we performed the assay in the presence of ethylene glycol-bis(β-aminoethyl ether)-N,N,NL,NL-tetraacetic acid (EGTA), a Ca^2+^ chelator (Fig. 2A). In contrast with the addition of Ca^2+^, the addition of EGTA completely mitigated the inhibitory effect of His-CaM on TSC2-Rheb interaction. To further confirm the role of Ca^2+^/CaM in TSC2-Rheb binding, we used a Ca^2+^-binding defective CaM mutant (DA) with mutations D21A, D57A, D94A, and D130A (34). The addition of wild-type CaM (WT) significantly reduced TSC2-Rheb binding in a dose-dependent manner. In contrast, the DA mutant did not interfere with the binding (Fig. 2B). The addition of either WT and DA did not affect the protein levels of GFP-TSC2 or HiBiT-Rheb during these assays, as the levels of these proteins remained consistent (Fig. 2C), ruling out the possibility that the observed reduction in HiBiT-Rheb in the IP fraction was attributed to a decrease in total HiBiT-Rheb rather than its binding to TSC2. These findings strongly suggest that CaM prevents the TSC2-Rheb interaction in a Ca^2+^-dependent manner (Fig. 2D).

**Figure 2.**
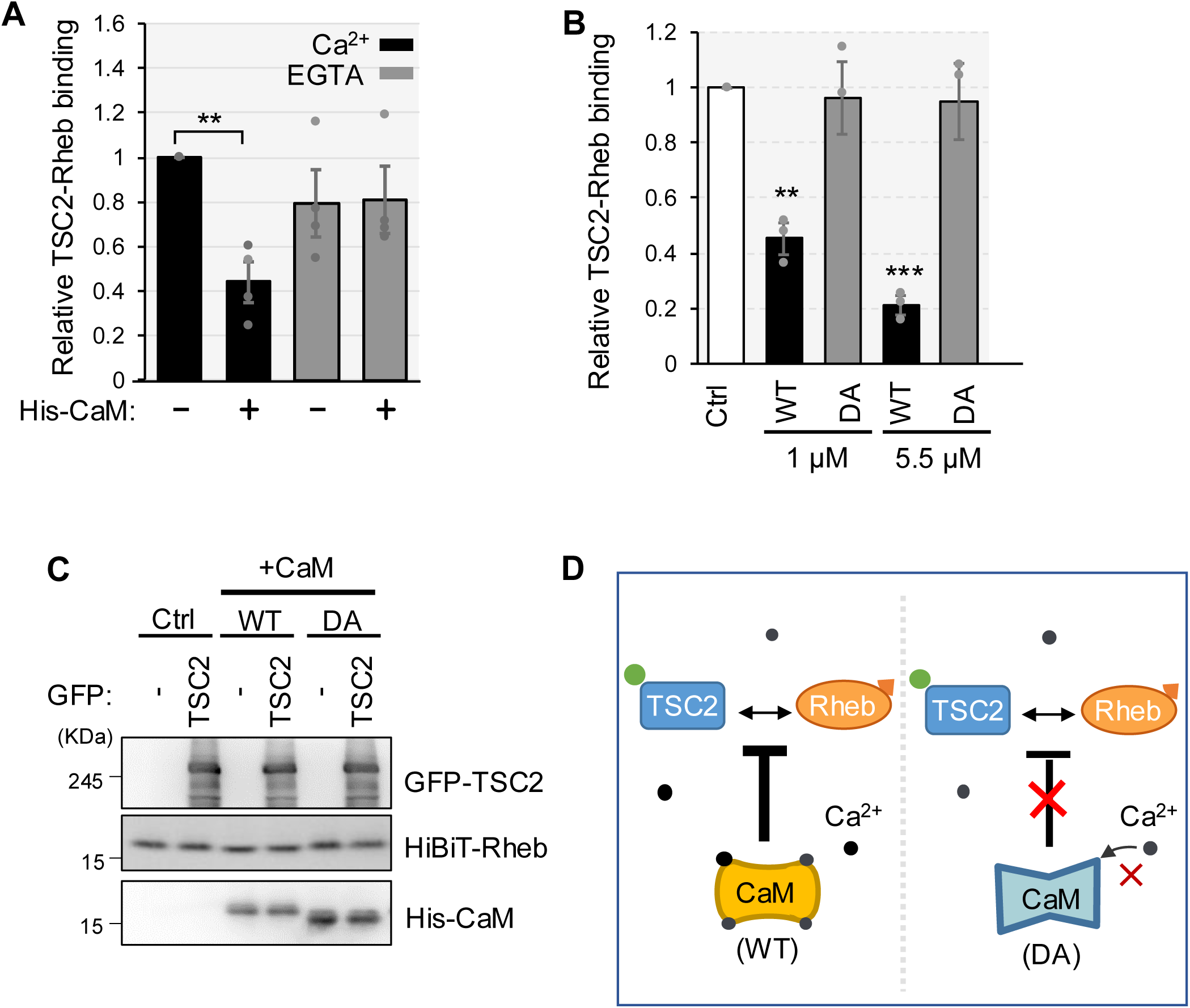
Ca^2+^ is required for CaM to prevent TSC2-Rheb interaction. (A) HEK293T cells were transiently transfected with the plasmid encoding Strep-GFP-TSC2 or HiBiT-Rheb. After 24 h of transfection, cells were lysed and respective cell lysates were mixed in the absence (-) or presence (+) of purified His-CaM (1 μM) together with 100 μM of CaCl_2_ (Ca^2+^) or 5 mM EGTA (EGTA). Subsequent procedures were done as in Fig 1(C). Graphs represent mean ± SEM of four independent experiments, One-way ANOVA with Tukey’s test, **p <0.01. (B) HEK293T cells were transiently transfected with the plasmid encoding Strep-GFP-TSC2 or HiBiT-Rheb. After 24 h of transfection, cells were lysed and respective cell lysates were mixed in the absence (Ctrl) or presence of purified 1 μM or 5.5 μM of wild-type His-CaM (WT) or Ca^2+^ binding-defective His-CaM (DA) together with 100 μM of CaCl_2_. Subsequent procedures were done as in (A). Graphs represent mean ± SEM of three independent experiments, One-way ANOVA with Tukey’s test, **p <0.01, ***p <0.001. (C) The (B) samples were also analyzed with western blotting using the anti-GFP and anti-6×His antibody and LgBiT blotting for detecting HiBiT-Rheb. (D) Schematic diagram of (B). WT CaM prevents TSC2 and Rheb interaction, while Ca^2+^ binding-defective CaM mutant (DA) does not interfere with the interaction.

### CaM binding to TSC2 reduces the ability of TSC2 to interact with Rheb

To further substantiate the hypothesis that CaM binding to TSC2 triggers to prevent TSC2-Rheb interaction, we used a TSC2 mutant lacking the CaM-binding region (residues 1717–1732) (TSC2ΔCaM) in the HiBiT lytic assay. GFP-TSC2ΔCaM exhibited reduced Rheb binding than that of wild-type GFP-TSC2 (Fig. 3A), consistent with cryo-EM analysis predictions that residues 1716–1731 of the TSC2 GAP domain support TSC2–Rheb interaction (21). Furthermore, compared with TSC2 WT, the TSC2ΔCaM mutant was less susceptible to the reduction in Rheb binding induced by His-CaM (Fig. 3B); however, the addition of His-CaM resulted in some residual reduction in TSC2ΔCaM-Rheb interaction, likely owing to residual binding of purified His-CaM to TSC2ΔCaM. Indeed, IP with an anti-His antibody indicated reduced, but not completely abolished, His-CaM binding to GFP-TSC2ΔCaM compared with its binding to wild-type GFP-TSC2 in this experimental setting (Fig. 3C and D). These results suggest that CaM prevents TSC2-Rheb interactions, at least partially, through its binding to the TSC2 GAP domain.

**Figure 3.**
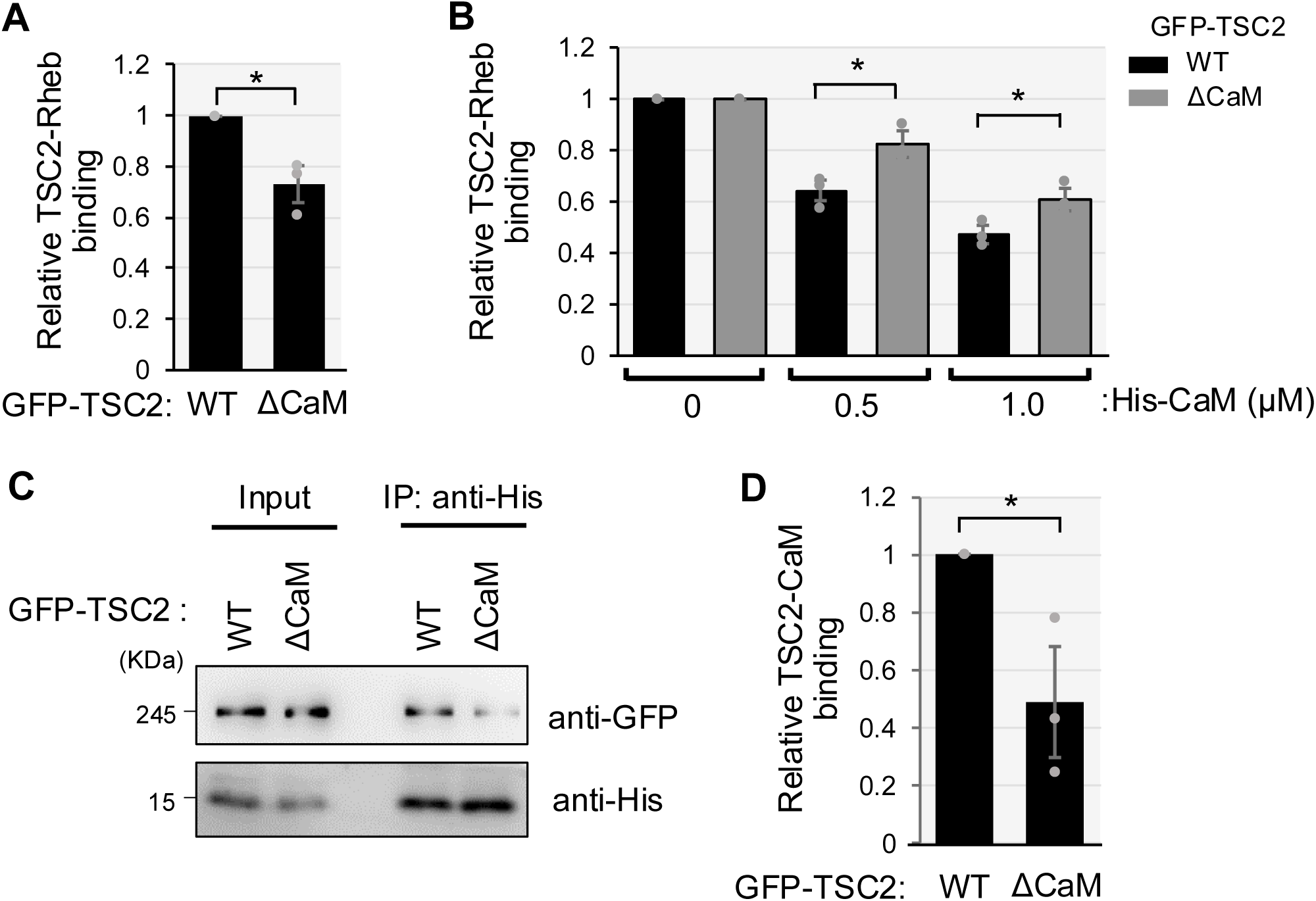
Binding of CaM to TSC2 contributes to the attenuation of TSC2-Rheb interaction. (A) HEK293T TSC2 KO cells were transiently transfected with the plasmid encoding wild-type (WT), CaM binding region deleted mutant (ΔCaM) of Strep-GFP-TSC2(GFP-TSC2) or HiBiT-Rheb. After 24 h of transfection, cells were lysed and respective cell lysates were mixed. Then, the cell lysates were subjected to immunoprecipitation using the anti-GFP antibody for overnight, followed by the addition of Protein G magnetic beads. After washing the beads, the amounts of HiBiT and GFP were monitored as described in Experimental Procedures. Graphs represent mean ± SEM of three independent experiments, Two-tailed unpaired Student’s t-test, *p <0.05. (B) HEK293T TSC2 KO cells were transiently transfected with the plasmid encoding wild-type (WT), CaM binding region deleted mutant (ΔCaM) of Strep-GFP-TSC2(GFP-TSC2) or HiBiT-Rheb. After 24 h of transfection, cells were lysed and respective cell lysates were mixed in the absence (0 μM) or presence (0.5 μM, 1.0 μM) of purified His-CaM together with 100 μM of CaCl_2_. Subsequent procedures were done as in (A). Graphs represent mean ± SEM of three independent experiments, Two-tailed unpaired Student’s t-test, *p <0.05. (C) HEK293T TSC2 KO cells were transiently transfected with the plasmid encoding wild-type (WT), CaM binding region deleted mutant (ΔCaM) of Strep-GFP-TSC2 (GFP-TSC2). Cell lysates were subjected to immunoprecipitation (IP) using anti-6×His antibody following His-CaM addition and analyzed by Western blotting with anti-GFP and anti-6×His antibodies. (D) Quantitation of the relative intensity of GFP-TSC2 to His-CaM of (C). Graphs represent mean ± SEM of three independent experiments.

### CaM prevents TSC2 binding to Rheb without affecting TSC complex assembly

TSC1 forms a TSC complex with TSC2 and TBC1D7, contributing to the stabilization of TSC2 and its GAP activity (18, 19, 22). Several studies have suggested that insulin signaling prevents the function of the TSC complex by inducing its disassembly (35, 36). Besides, a recent report has reported that TSC1 binds to the lysosomal membrane via PI(3,5)P_2_ and is required to translocate the TSC complex to the lysosome (37). Therefore, TSC1 is a key player that regulates the TSC complex, together with TSC2. We investigated whether CaM binding-mediated inhibition of the TSC2-Rheb interaction was affected by a change in the interaction between TSC1 and TSC2.

To evaluate the effect of CaM addition on TSC1–TSC2 binding and TSC2–Rheb binding simultaneously, myc-tagged TSC1 (TSC1-myc) was co-expressed with HiBiT-Rheb and subjected to a TSC2-Rheb binding assay by mixing with cell lysates containing GFP-TSC2. After purification of GFP-TSC2 immunoprecipitates in the presence of His-CaM (WT) or His-CaM (DA), bound TSC1-myc was detected using western blotting (Fig. 4A and B), and bound HiBiT-Rheb was detected using the assay system (Fig. 4C). The binding between TSC1-myc and GFP-TSC2 was not significantly affected by the addition of either His-CaM (WT) or His-CaM (DA) (Fig. 4A and B), whereas the addition of His-CaM (WT), but not His-CaM (DA), consistently decreased the formation of the TSC2-Rheb complex (Fig. 4C). These results suggest that CaM binding to TSC2 does not affect TSC1–TSC2 complex formation, and the CaM-mediated reduction of TSC2-Rheb interaction is not mediated by the change of TSC1-TSC2 complex.

**Figure 4.**
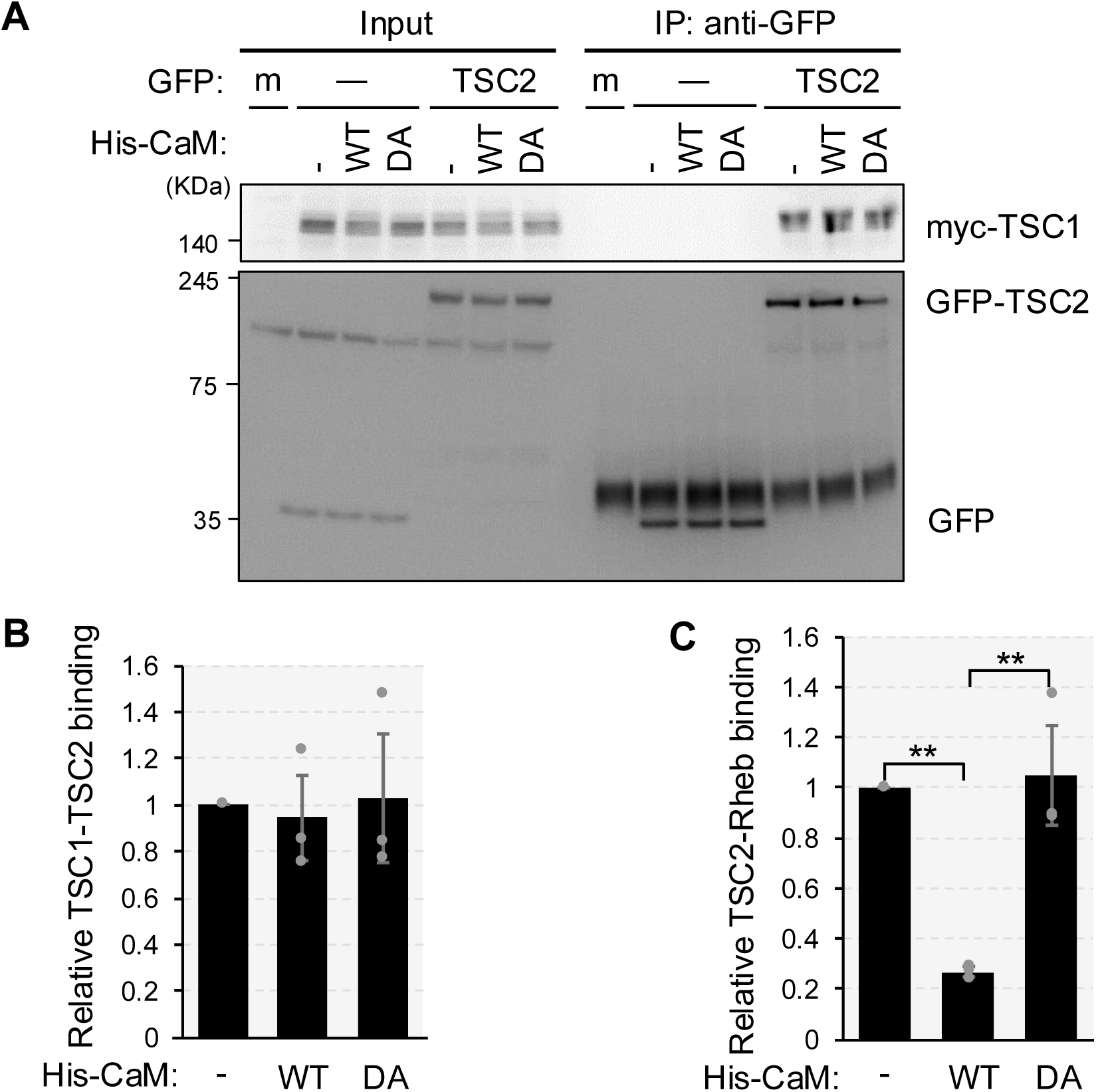
Ca^2+^/CaM alters TSC2-Rheb binding without affecting TSC complex assembly. (A) HEK293T cells were mock transfected (m) or transiently transfected with the plasmid encoding Strep-GFP (GFP, -), Strep-GFP-TSC2 (GFP-TSC2) or with both the plasmid encoding HiBiT-Rheb and the plasmid for TSC1-myc. After 24 h of transfection, cells were lysed and respective cell lysates were mixed in the absence (-) or presence of purified wild-type (WT) or Ca^2+^ binding-defective mutant (DA) of His-CaM (5.5 μM) together with 100 μM of CaCl_2_. Then, the cell lysates were subjected to immunoprecipitation (IP) using the anti-GFP antibody and analyzed by Western blotting with myc and anti-GFP antibodies. (B) Quantitation of the relative intensity of GFP-TSC2 to TSC1-myc of (A). Graphs represent mean ± SEM of three independent experiments. (C) The samples in (A) were also analyzed by HiBiT lytic assay as described in Experimental Procedures. Graphs represent mean ± SEM of three independent experiments, One-way ANOVA with Tukey’s test, **p <0.01.

### Increase in intracellular Ca^2+^ causes CaM binding to the region of TSC2 in cells

To investigate the binding of CaM to TSC2 in response to Ca^2+^ mobilization, we used R-GECO1 (38). R-GECO1 contains a circularly permutated mApple (cp-mApple) linked to CaM and an M13 peptide derived from myosin light chain kinase at its C- and N-terminus, respectively (Fig. 5A). An increase in intracellular Ca^2+^ concentration enhances cp-mApple fluorescence owing to the binding of CaM to the M13 peptide. We replaced the M13 peptide with a 16 amino acid peptide corresponding to the CaM-binding region of TSC2 (residues 1717–1732) to create R-GECO1-TSC2 WT (Fig. 5A) and examined its sensitivity to changes in intracellular Ca^2+^. We also constructed a mutant version of R-GECO1-TSC2 (WQLQ), with mutations W1717Q and W1727Q in the CaM-binding region of TSC2 that lost the ability to bind CaM (data not shown). We expressed these R-GECO1 variants in HeLa cells, added ionomycin to promote Ca^2+^ influx, and monitored changes in the fluorescence signal. Cells expressing R-GECO1-TSC2 WT exhibited increased fluorescence with ionomycin, different from those expressing R-GECO1-TSC2 (WQLQ) (Fig. 5B and movies S1–S3), although the dynamic range and/or duration of the signal was lower than that of R-GECO1. These different responses might reflect differences in affinity between the M13 and TSC2 peptides to CaM. Collectively, these results suggest that CaM binds to TSC2 in response to Ca^2+^ mobilization in cells.

**Figure 5.**
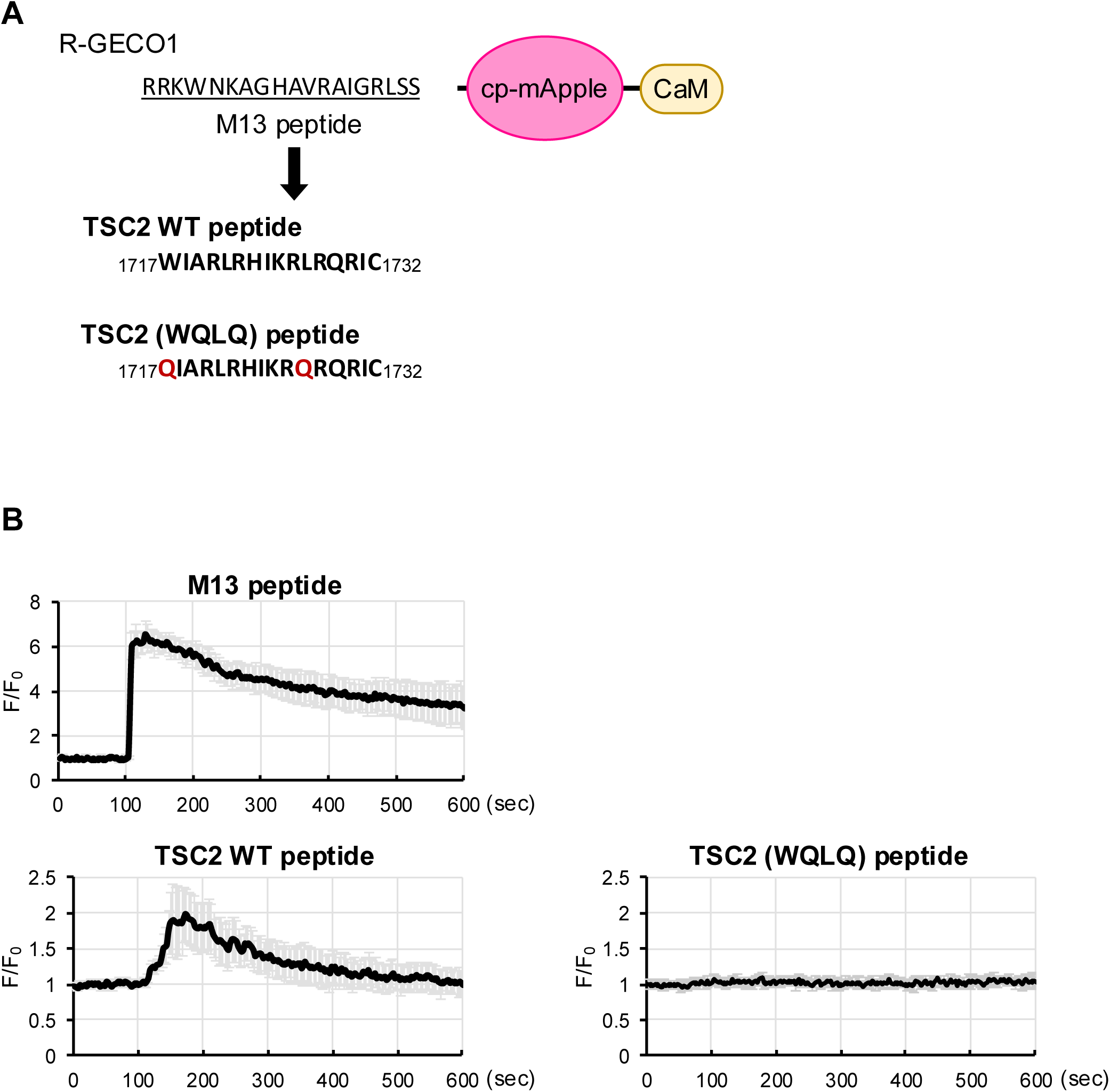
CaM binds the GAP domain of TSC2 in a Ca^2+^-dependent manner in cells. (A) Schematic representation of CaM binding peptide substituted R-GECO1 construct. The M13 peptide (CaM binding peptide from myosin light chain) was replaced with the CaM binding region of TSC2 (residues 1717-1732) or CaM binding defective mutation (red) of TSC2 peptide (WQLQ). (B) HeLa cells were transiently transfected with the plasmid encoding R-GECO1(M13 peptide), R-GECO1(TSC2 WT peptide) or R-GECO1(TSC2 (WQLQ) peptide). The medium was replaced with HBSS for 15 min, and then ionomycin (2.5 μM) was added at 105 s. The ratio of fluorescence signals for R-GECO1 to the average signals before ionomycin stimulation (F_0_) was shown. Graphs are shown as mean ± SD (n = 15).

### Intracellular Ca^2+^ mobilization alters mTORC1 activity via CaM

We previously demonstrated that the addition of amino acids to amino acid-starved cells induces Ca^2+^ entry from the extracellular medium, leading to mTORC1 activation (16). To further examine whether Ca^2+^ mobilization modulates mTORC1 activity, we evaluated the effect of carbachol on the phosphorylation of ribosomal protein S6 kinase 1 (S6K1), a readout of mTORC1 activity. Carbachol is a cholinergic agonist that increases cytosolic Ca^2+^ levels by inducing the release of Ca^2+^ from the endoplasmic reticulum and intracellular Ca^2+^ stores (39). In HEK293 cells, the addition of carbachol to the cells starved of serum for 3 h led to an increase in the phosphorylation of Thr389 on S6K1 (Fig. 6A). Using R-GECO1, we also observed that carbachol induced a transient increase in intracellular Ca^2+^ concentration (Fig. 6B, movie S4). Furthermore, pretreatment with BAPTA-AM, an intracellular Ca^2+^ chelator, completely abolished carbachol-induced phosphorylation of S6K1. This indicates that an increase in Ca^2+^ levels triggered by carbachol promotes S6K1 phosphorylation (Fig. 6C and D).

**Figure 6.**
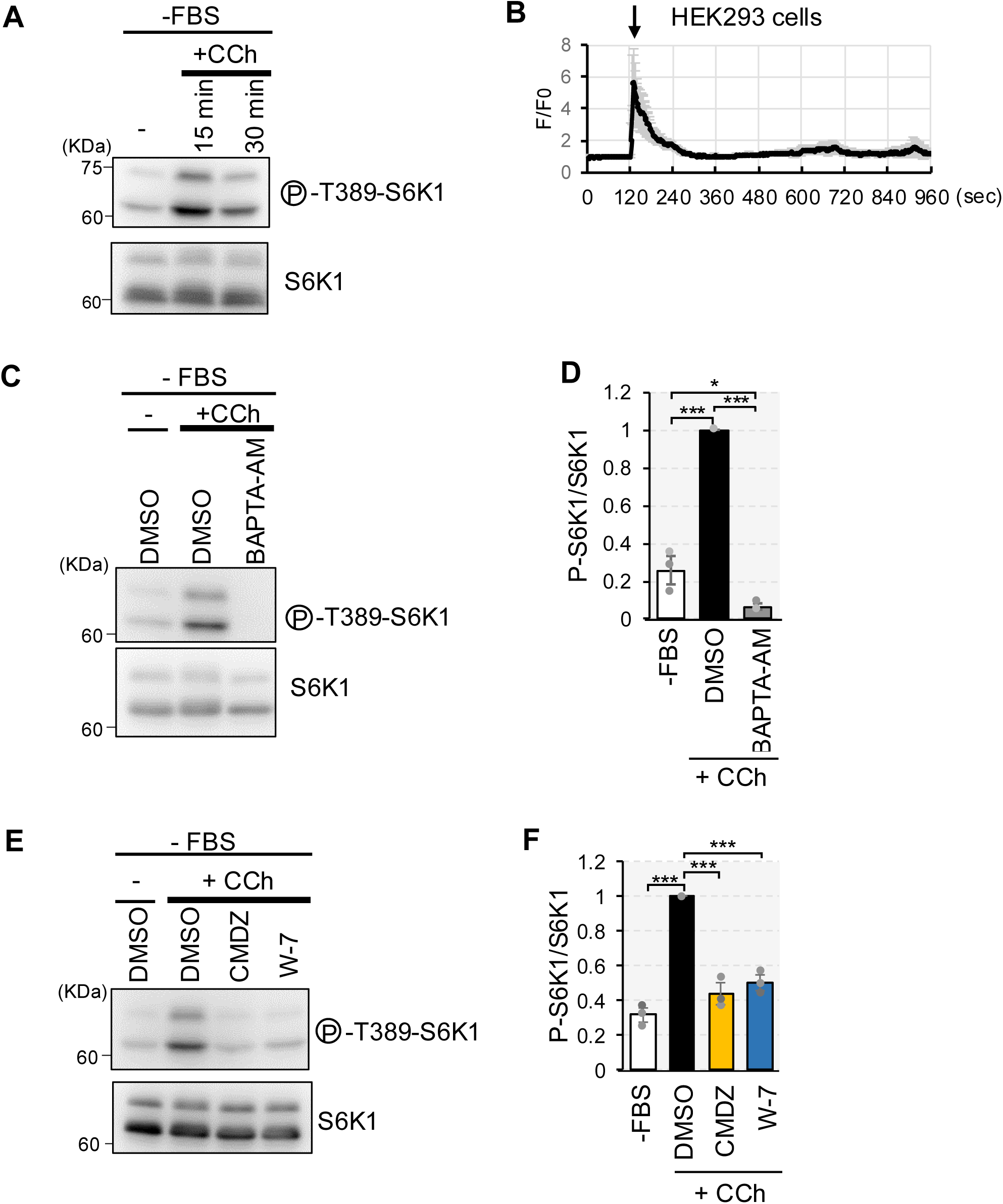
The intracellular Ca^2+^ rise signals to mTORC1 via CaM. (A) HEK293 cells were cultured in serum-starved condition (−FBS) for 3 h and then stimulated with or without (-) carbachol (100 μM) for 15 or 30 min (+CCh). Cell lysates were analyzed by Western blotting with the indicated antibodies. (B) HEK293 cells transiently transfected with the plasmid encoding R-GECO1 were deprived of serum for 3 h, then carbachol (100 μM) were added at 120 s. Ratio of fluorescence signals for R-GECO1 to the average signals before carbachol stimulation (F_0_) were shown. Graphs are shown as mean ± SD (n = 12). (C) HEK293 cells were cultured in serum-starved condition (−FBS) for 3 h pretreated with DMSO or BAPTA-AM (50 µM) for the last 1 h and then stimulated with or without (-) carbachol (100 μM) for 15min (+CCh). Cell lysates were analyzed by Western blotting with the indicated antibodies. (D) Quantitation of the relative intensity of phospho-T389-S6K1 to total S6K1 of (C), in the carbachol stimulation (DMSO) was set to 1. Graphs represent mean ± SEM of three independent experiments, One-way ANOVA with Tukey’s test, * p < 0.05, *** p < 0.001. (E) HEK293 cells were cultured in serum-starved condition (−FBS) for 3 h pretreated with DMSO or CaM inhibitors (W-7 (30 µM) or Calmidazolium (CMDZ, 30 μM)) for the last 15 min (CMDZ) or 5 min (W-7) and then stimulated with or without (-) carbachol (100 μM) for 15min (+CCh). Cell lysates were analyzed by Western blotting with the indicated antibodies. (F) Quantitation of the relative intensity of phospho-T389-S6K1 to total S6K1 of (E), in the carbachol stimulation (DMSO) was set to 1. Graphs represent mean ± SEM of three independent experiments, One-way ANOVA with Tukey’s test, *** p < 0.001.

To further examine whether CaM was also involved in carbachol-induced mTORC1 activation, cells were pretreated with different CaM inhibitors, W-7 or Calmidazolium (CMDZ), before carbachol treatment. The CaM inhibitors suppressed the carbachol-induced increase in S6K1 phosphorylation (Fig. 6E and F). These results suggest that intracellular Ca^2+^ mobilization promotes mTORC1 activation via CaM.

Akt-mediated TSC2 phosphorylation controls the dissociation of the TSC complex from the lysosomal membrane, leading to Akt-dependent mTORC1 activation via TSC2 (11, 25). To examine whether Ca^2+^/CaM affects the phosphorylation status of TSC2, we treated cells with carbachol with or without pretreatment with BAPTA-AM or W-7. Treatment with carbachol did not increase the phosphorylation of TSC2 at Thr1462, whereas insulin significantly increased the phosphorylation (Fig. 7A, B, and Fig. S1A, B). Furthermore, pretreatment with BAPTA-AM or W-7 had no impact on TSC2 phosphorylation, even though the phosphorylation of S6K1 was repressed by these pretreatments (Fig. 7A and B).

**Figure 7.**
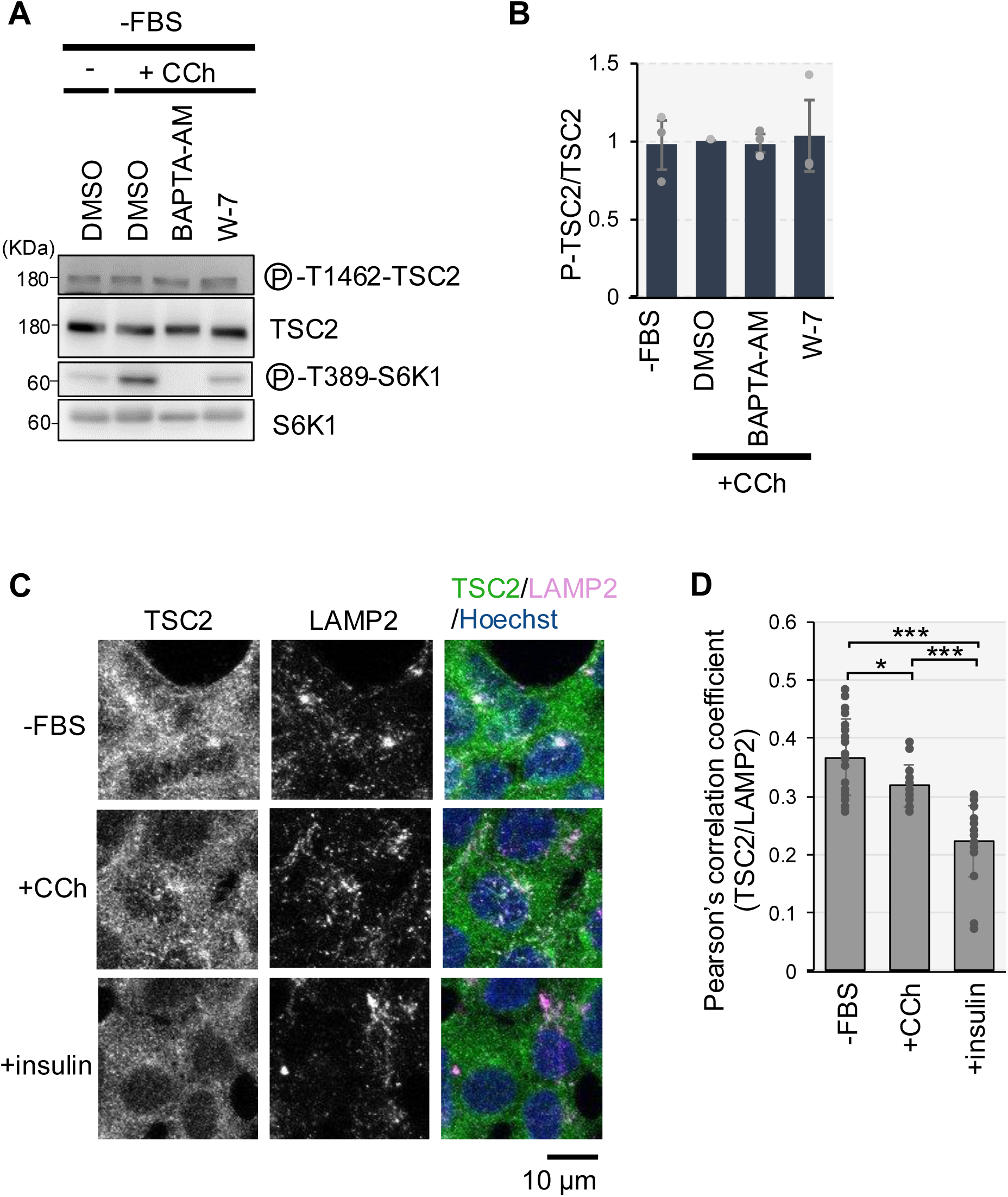
Ca^2+^/CaM affects TSC2 localization without altering TSC2 phosphorylation. (A) HEK293 cells were starved of serum (−FBS) for 3 h, pretreated with DMSO or BAPTA-AM (50 µM) or W-7 (30 μM) for the last 1 h (BAPTA-AM) or 5 min (W-7) and then stimulated with or without (-) carbachol (100 μM) for 15 min (+CCh). Cell lysates were analyzed by Western blotting with the indicated antibodies. (B) Quantitation of the relative intensity of phospho-T1462-TSC2 to total TSC2 of (A), in which the carbachol stimulation (DMSO) was set to 1. Graphs represent mean ± SEM of three independent experiments. One-way ANOVA with Tukey’s test. (C) HEK293 cells were starved of serum (-FBS) for 3 h, and then treated with carbachol (100 μM) or insulin (100 nM) for 15 min. The cells were then immunostained with anti-TSC2 and anti-LAMP2 antibodies. Merged images of TSC2 (green), LAMP2 (magenta) and nuclei staining with Hoechst 33342 (blue) are also shown. Scale bar, 10 µm. (D) Colocalizations of TSC2 with LAMP2 were quantified, and Pearson’s correlation coefficient is shown. n = 20. One-way ANOVA with Tukey’s test, * p < 0.05, *** p < 0.001.

We also investigated whether carbachol treatment affected the subcellular localization of TSC2. Consistent with previous reports (11), a subpopulation of TSC2 co-localized with the lysosomal marker LAMP2 under serum-starved conditions, whereas treatment with insulin induced the dissociation of TSC2 from lysosomes marked with LAMP2 (Fig. 7C and D). Notably, carbachol treatment also induced the dissociation of TSC2 from the lysosomes, albeit to a lesser extent than insulin treatment (Fig. 7C and D). Collectively, these results suggest that Ca^2+^/CaM primarily affects TSC2-Rheb interaction to modulate mTORC1 activity.

## Discussion

We previously reported that extracellular Ca^2+^ influx induced by amino acids is essential for mTORC1 activation, mediated by the ubiquitous Ca^2+^ sensor CaM via the TSC2-Rheb axis (16). In this study, we elucidate the underlying molecular mechanism, demonstrating that Ca^2+^/CaM inhibits TSC2 binding to Rheb, thereby driving mTORC1 activation. The CaM-binding region within TSC2 is proposed to be a critical α-helix for TSC2-Rheb interaction (Fig. 1A) (21). Consistent with this model, the TSC2 mutant lacking this α-helix corresponding to the CaM-binding region exhibited reduced binding to Rheb (Fig. 3A). We further investigated the association between Ca^2+^ signaling and mTORC1 activation. Treatment with carbachol increased Ca^2+^ levels and mTORC1-dependent S6K1 phosphorylation, whereas pretreatment with CaM inhibitors suppressed this phosphorylation (Fig. 6), suggesting that Ca^2+^/CaM plays a crucial role in mTORC1 activation. It has been reported that carbachol-induced S6K1 phosphorylation was independent of the PI3K–Akt pathway (40, 41). Consistent with these reports, TSC2 phosphorylation at Thr1462 remained unaffected by carbachol or CaM inhibitors (Fig. 7A and B). Thus, CaM appears to mediate carbachol-induced Ca^2+^-dependent mTORC1 activation by directly acting on TSC2 rather than affecting upstream factors.

Our findings reveal a novel regulatory mechanism of TSC2 action on Rheb via Ca^2+/^CaM. The TSC complex acts as a potent negative regulator of mTORC1 through its action on Rheb. Recent studies have highlighted that changes in TSC2 subcellular localization, rather than inhibition of its GAP activity, are crucial for regulating Rheb activity (11, 15, 25). During serum or amino acid starvation, TSC2 relocates from the cytoplasm to the lysosomal surface, where it likely inactivates lysosome-localized Rheb. Conversely, growth factors such as insulin induce the dissociation of TSC2 from lysosomes, leading to the reversal of Rheb suppression. These growth factors activate Akt, which in turn directly phosphorylates and inactivates TSC2. The phosphorylation of TSC2 appears to be a key mechanism in controlling the lysosomal localization of TSC2 and/or its turnover at the lysosomal surface (11, 25). Earlier studies have also suggested other potential mechanisms by which Akt-mediated phosphorylation of TSC2 disrupts the TSC complex (35, 36, 42) and promotes the degradation of TSC2 (43). It was also suggested that Ca^2+^/CaM could cause the translocation of TSC2 to the nucleus by promoting its dissociation from the membrane (44). However, our current findings indicate that Ca^2+^/CaM-mediated regulation of TSC2 does not significantly affect Akt-dependent phosphorylation of TSC2 or TSC complex integrity (Figs. 4 and 7A). Notably, we observed that the localization of the TSC2 subpopulation changed upon carbachol treatment (Fig. 7C and D). The lysosomal localization of TSC2 under serum starvation depends on Rheb, and Akt-mediated TSC2 phosphorylation can induce its dissociation from lysosomes (11). Therefore, the dissociation observed upon carbachol treatment is likely a consequence of disrupted TSC2-Rheb binding induced by Ca^2+^/CaM binding to TSC2. Collectively, our study suggests that by acting as a competitor of Rheb, Ca^2+^/CaM can prevent the action of TSC2, leading to Rheb activation and subsequent mTORC1 activation (Fig. 8).

**Figure 8.**
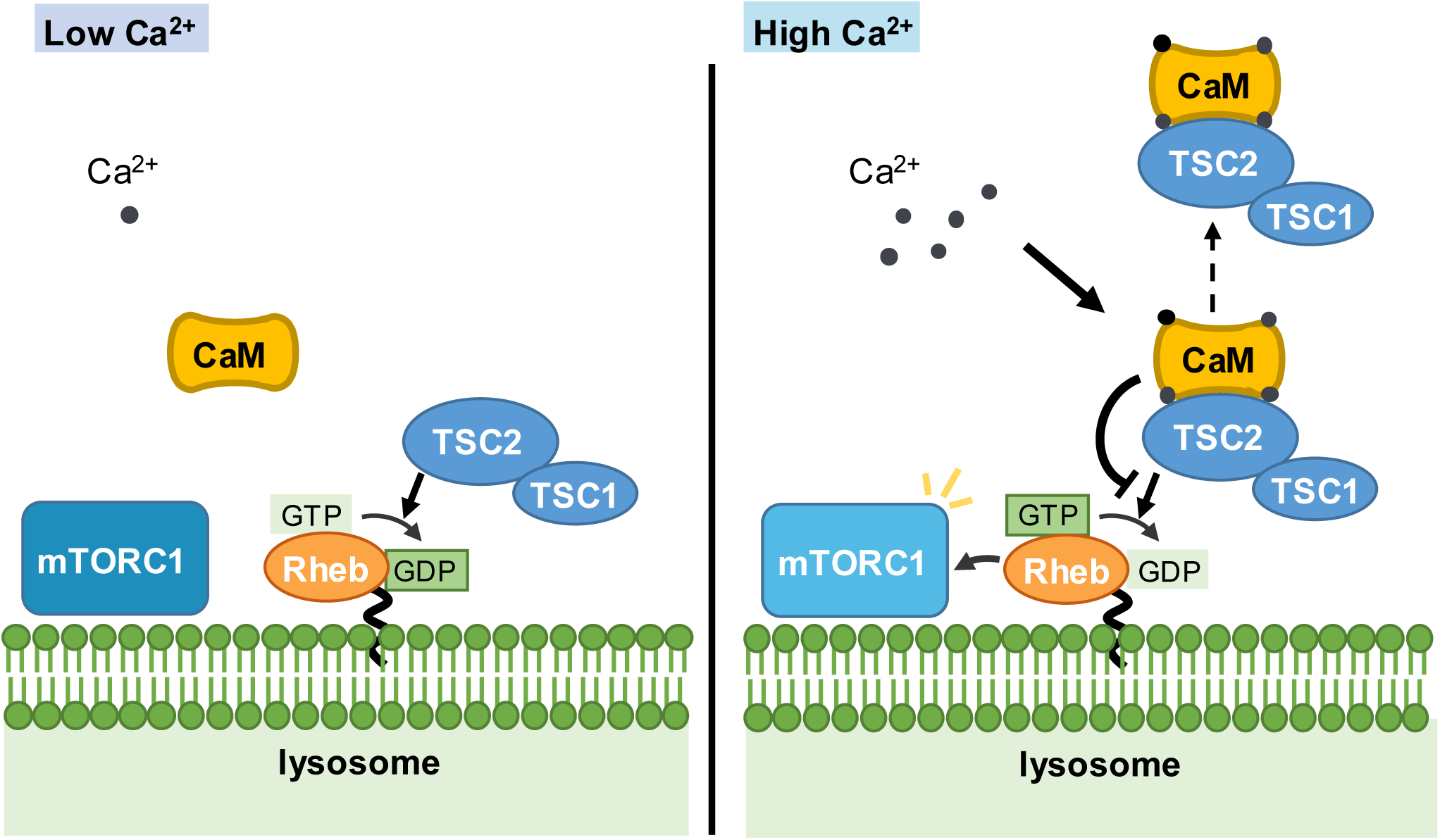
A proposed model for Ca^2+^/CaM regulation of mTORC1 via TSC2-Rheb. In the low Ca^2+^ condition, most of CaM is not fully occupied by Ca^2+^, and TSC2 can bind and inactivate Rheb by GAP activity of TSC2. In the high Ca^2+^ condition, it facilitates CaM binding to TSC2. CaM binding to TSC2 inhibits the interaction between TSC2 and Rheb without affecting TSC complex integrity, and results in the dissociation of TSC2 from the lysosome where Rheb resides. Thus, Ca^2+^/CaM facilitates mTORC1 activation by preventing TSC2 from inactivating Rheb.

We estimated the IC_50_ of CaM inhibiting the TSC2-Rheb interaction to be approximately 1.1 μM (Fig. 1E). Previously, Noonan et al. estimated the K_d_ of the CaM-TSC2 peptide as approximately 1 μM (32), although a link between the binding of CaM to TSC2 and mTORC1 signaling remained unexplored. The similar K_d_ and IC_50_ values further suggest that direct binding of CaM to TSC2 hinders TSC2 from binding to Rheb. CaM contains over 300 target peptides and is involved in various essential biological processes through binding target proteins (45). Most target proteins have a high affinity for CaM (K_d_ in the nM range). However, some targets such as caldesmon, B50 (neuromodulin), and Rab3A show low affinity (K_d_ in µM range, caldesmon: 0.8 µM, B50: 4.2 µM, and Rab3A: 18–22 µM) (46–48). Our analysis using the R-GECO1-based system (Fig. 5) indicated that TSC2–CaM binding occurred in response to Ca^2+^ mobilization in cells, demonstrating that TSC2 is a CaM target protein. The relatively low affinity of TSC2 for CaM may be beneficial in preventing excessive suppression of TSC2 by CaM under basal conditions. This prevents potential adverse effects on normal cell growth owing to enhanced Rheb and subsequent mTORC1 activation. This prompt switching between on- and off-like states with Ca^2+^ signaling allows the mTORC1 pathway to respond appropriately to upstream stimuli.

CaM is a ubiquitous protein with numerous binding partners. However, the mechanisms determining CaM selectivity for its target proteins remain unclear. CaM binding partners share several common characteristics: 1) an α-helical conformation; 2) a hydrophobic motif with two or three hydrophobic amino acids repeated at specific intervals (such as 1-10 motif, 1-14 motif, or 1-16 motif); and 3) a positive charge owing to multiple basic amino acids (49). The CaM target sequence of TSC2 fits this pattern, forming an α-helix and matching the 1-10-14 motif (1718I, 1727L, 1731I). The combination of required Ca^2+^ levels and target sequences of CaM likely affects its selectivity.

We previously showed that CaM inhibitors, such as W-7 and CMDZ, can prevent the binding of CaM to TSC2 and suppress mTORC1 signaling (16). However, such inhibitors also prevent Ca^2+^/CaM binding to other target proteins, limiting their therapeutic potential owing to a lack of selectivity. Instead, small molecules that specifically target the CaM-TSC2 interaction present a more promising approach. Our findings reveal a new molecular mechanism of Ca^2+^/CaM-mediated regulation of TSC2 and identify potential drug targets that may block the binding of CaM to TSC2, thereby enhancing the inhibitory action of TSC2 on Rheb and ultimately controlling mTORC1 activity.

## Experimental procedures

### Antibodies and Reagents

Anti-pThr389-S6K1 (#9234), anti-S6K1 (49D7, #2708), anti-pThr1462-TSC2 (5B12, #3617), anti-TSC2 (D93F12, #4308) were purchased from Cell Signaling Technology (Danvers, MA, USA). Anti-Myc (9E10, sc-40), anti-GFP (B-2, sc-9996) were from Santa Cruz Biotechnology (Dallas, TX, USA). Anti-6xHis tag (66005-1-Ig) and anti-Strep II tag (NBP2-41076) were from Proteintech (Rosemont, IL, USA) and Novus Biologicals (Centennial, CO, USA), respectively. Rabbit polyclonal antibody (pAb) against GFP was prepared previously (33). Horseradish peroxidase (HRP)-conjugated goat antibodies against mouse IgG and rabbit IgG were from Jackson ImmunoResearch (West Grove, PA, USA). For immunofluorescence, TSC2 (D93F12, Cell Signaling Technology #4308) and LAMP2 (H4B4, Proteintech 65053-1-Ig) were used as primary antibodies. AlexaFluor488-conjugated donkey anti-rabbit IgG and AlexaFluor555-conjugated donkey anti-mouse IgG secondary antibodies were purchased from Invitrogen (Carlsbad, CA, USA). Carbachol (sc-202092) was purchased from Santa Cruz Biotechnology (Dallas, TX, USA). BAPTA-AM solution (348–05451) was purchased from Wako (Tokyo, Japan). Calmidazolium (14442) and Ionomycin (10004974) were from Cayman Chemical (Ann Arbor, MI, USA), and W-7 hydrochloride (A2409) was from TCI (Tokyo, Japan). Hoechst 33342 (04929–82) was purchased from Nacalai tesque (Kyoto,Japan).

### Plasmids

The expression plasmid for R-GECO1 (CMV-R-GECO1) was a gift from Robert Campbell (Addgene plasmid #32444) (38). pcDNA3.1-myc-TSC1 was a gift from Cheryl Walker (Addgene plasmid #12133) (42). pStrepHA-SGFP2 was prepared previously (33). The expression plasmid for HiBiT-Rheb (pEXPR-HiBiT-Rheb) was generated by inserting annealed oligos (ctagcgtgagcggctggcggctgttcaagaagattagcgggagctccggaccggttggctcgg and gatcccgagccaaccggtccggagctcccgctaatcttcttgaacagccgccagccgctcacg) into NheI and BamHI sites of pEXPR-IBA105-B-Rheb (16).

The expression plasmid for GFP-TSC2 (pSGFP2-C3-TSC2) was generated by inserting the TSC2-containing fragment from the digestion of pcDNA3-FLAG-TSC2 (addgene #14129) by BamHI into the BamHI site of pSGFP2-C3. The expression plasmid for Strep-HA-GFP-TSC2 (pStrepHAGFP-TSC2), the AgeI-HpaI fragment from pSGFP2-C3-TSC2 was inserted into AgeI and PmeI sites of pStrepHA-SGFP2. To generate an expression plasmid for GFP-TSC2Δ1717–1732 (ΔCaM), a PCR fragment having the corresponding deletion was amplified using primers (gctccaaccccaccgatatctacccttcgaaggaggaagccgcctactcc and aagctgcaataaacaagttaacaacaacaattgcattca) and inserted into HpaI and EcoRV sites. Note that these TSC2 plasmids encodes 1784 residues that correspond to isoform 4 under Uniprot, and we used this numbering throughout this paper except for the antibody name (the anti-pThr1462-TSC2 antibody).

To generate the plasmid for 6xHis-tagged CaM (pET24His-CALM2), the CALM2 fragment was amplified by PCR using primers (catcatcatcatggggatcccgctgaccaactgactgaaga and tggtggtggtggtgctcgagtcactttgctgtcatcattt) and pGEX4T-CALM2 (lab stock) as a template, and inserted into BamHI and XhoI sites of the pET24His plasmid. The plasmid for N-terminally His-tagged CaM DA (pET24His-CALM2 D21A, D57A, D94A, D130A) was generated by insering the codon-optimized synthetic gene fragment (ThermoFisher, Strings DNA Fragments) into BamHI amd XhoI sites of pET24His plasmid.<colcnt=3>

To generate plasmids for CMV-R-GECO1-TSC2(1717–1732) and CMV-R-GECO1-TSC2(1717–1732)(W1717Q,1727Q), inverse PCR was performed using CMV-R-GECO1 as a template and primers set (cgccacatcaagcggctccgccagcggatctgccccgtggtttccgagcggat and gagccgcttgatgtggcggagccgggcaatccatgatgagtcgaccatggtgg) and (cgccacatcaagcggcagcgccagcggatctgccccgtggtttccgagcggat and ctgccgcttgatgtggcggagccgggcaatctgtgatgagtcgaccatggtgg), respectively. Amplified fragments were assembled by using NEBuilder®HiFi DNA Assembly Master Mix (New England Biolabs, Ipswich, MA). All DNA constructs were verified by DNA sequencing.

### Mammalian Cell Culture and Transfection

HEK293, HEK293T, and HeLa cells were grown at 37°C in Dulbecco’s modified Eagle’s medium (DMEM) with 1,000 mg/L glucose (Shimadzu, Kyoto, Japan) or with 4,500 mg/L glucose (for HEK293T cells) supplemented with 5% fetal bovine serum under a 5% CO_2_ atmosphere. HEK293T TSC2KO cells were described previously (16). Transient transfection of the indicated plasmids was performed using Polyethylenimine “Max”(PEImax) (#24765, Polysciences Inc., Warrington, PA, USA) according to the manufacturer’s instructions.

For carbachol stimulation, HEK293 cells were washed once with serum-free DMEM and incubated in serum-free DMEM for 3 h. Carbachol were added to the final concentration of 100 µM following pretreatment with or without BAPTA-AM (50 µM), W-7 (30 µM) or calmidazolium (20 µM).

### Preparation of recombinant proteins

*Escherichia coli* Rosetta2 (DE3)pLysS was transformed with the plasmid, pET24His-CaM or pET24His-CaM D21A,D57A, D94A, D130A (DA), and cultured at 37°C 4 h after induction for expression with 1 mM isopropyl β-D-thiogalactopyranoside. Bacteria cells were lysed with lysis buffer His [20 mM Tris-HCl (pH 8.0), 300 mM NaCl, 5 mM imidazole, 1 mM phenylmethylsulfonyl fluoride, 5 mM benzamidine] and both His-tagged proteins were purified using TALON® Metal Affinity Resin from TAKARA (Clontech/Takarabio, Kusatsu, Shiga, Japan) and eluted by elution buffer [20 mM Tris-HCl (pH8.0), 200 mM NaCl, 150 mM imidazole]. The elution buffer was replaced with another buffer [40 mM HEPES-NaOH (pH7.4), 120 mM NaCl] for preparation to use for HiBiT-lytic assay.

### HiBiT Lytic Assay

HEK293T or HEK293T TSC2KO cells were transfected independently with each expression plasmids encoding SGFP2-TSC2 and TSC1-myc or HiBiT-Rheb. After 24 h, harvested cells were lysed at 4°C with lysis buffer HLA [40 mM HEPES-NaOH (pH7.4), 120 mM NaCl, 10 mM β-glycerophosphate, 10 mM MgCl_2_] containing 0.3% CHAPS and protease inhibitors (3 µg/mL leupeptin, 1 µM pepstatin A, 0.1 mM pefabloc). The cleared cell lysates were obtained by centrifugation at 13,000 × g for 10 min and mixed with each other (the lysate from cells expressing GFP-TSC2 and TSC1-myc and the lysate from cells expressing HiBiT-Rheb) to make them form complex. For Figure 4, the cell from cells expressing GFP-TSC2 and the lysate from cells expressing both TSC1-myc and HiBiT-Rheb were mixed. The purified recombinant His-CaM were added to the indicated final concentration, followed by adding CaCl_2_ (100 µM) or EGTA (5 mM). HiBiT lytic assay was performed as described previously (33) with slight modifications. In brief, the lysate was used for immunoprecipitation by incubation with rabbit anti-GFP pAb overnight followed by incubation with Protein G magnetic beads (Dynabeads® protein G, Thermo Fisher Scientific, Waltham, MA, USA) for 1 h. Collected magnetic beads were washed with buffer HLA containing CaCl_2_ or EGTA. After washing beads, the beads were suspended with passive lysis buffer (PLB) (Promega, Madison, WI, USA). Aliquots were taken and luciferase activities were measured using a Nano-Glo® Luciferase assay reagent kit (Promega) with a photon-counting type microplate reader (Monochromator Multimode Microplate Reader-Berthold Technologies Mithras2 LB 943, Bad Wildbad, Germany). For normalization of the amounts of immunoprecipitated Strep-SGFP2-fused proteins, activities of StrepTactin-conjugated alkaline phosphatase (Precision Protein StrepTactin-AP Conjugate, BIO-RAD, Hercules, CA, USA) were measured using a substrate contained in a Phospha-Light™ SEAP Reporter Gene Assay System (Thermo Fisher Scientific, Waltham, MA, USA) as described previously (33) or GFP fluorescence emission was measured using a microplate reader with excitation at 480 nm and emission at 520 nm (Infinite® 200 PRO, TECAN, Mannedorf, Switzerland).

### Cell Lysate Preparation, Pulldown and Western Blotting

Cells were washed once with ice-cold PBS(−) and lysed with lysis buffer TX [50 mM Tris-HCl (pH 7.5), 150 mM NaCl, 1% Triton X-100, 50 mM NaF, 10 mM β-glycerophosphate] containing 3 µg/mL leupeptin, 1 µM pepstatin A, and 0.1 mM pefabloc and centrifuged at 9000 × g for 10 min to obtain cell lysates. In Fig. 3C, HEK293T TSC2 KO cells were washed once with ice-cold PBS(−) and lysed with lysis buffer HLA. After centrifugation at 13000 × g for 10 min, the supernatant was incubated overnight with the anti-6×His-tag antibody following the addition of His-CaM. The immunocomplexes were collected by the addition of Dynabeads protein G (Invitrogen) and washed three times with lysis buffer HLA.

### Immunofluorescence

HEK293 cells (3 x10^5^ cells) were seeded into 35-mm dishes containing PLL-treated glass coverslips. After 24 h, cells were washed once with serum-free DMEM and incubated in serum-free DMEM for 3 h. Cells were left or treated with carbachol (100 µM) or insulin (100 nM) for 15 min. Cells were then washed once with PBS(-) and fixed with 4% paraformaldehyde in PBS(-) for 15 min, followed by washing twice with PBS(−). Then the cells were permeabilized with 0.1% Triton X-100/PBS(−) for 5 min, followed by washing twice with PBS(−). After blocking with 0.25% bovine serum albumin (BSA)/PBS(−) for 1 h, the cells were incubated with primary antibodies at 4°C overnight and washed five times with 0.25% BSA/PBS(−). The cells were then incubated with secondary antibodies at room temperature for 1 h in the dark and then washed five times with 0.25% BSA/PBS(−), stained with Hoechest 33342 (5 µM) for 10 min, and washed once with 0.25% BSA/PBS(−) before mounting. Images were acquired under an FV3000 confocal laser-scanning microscope equipped with a numerical aperture oil-immersion objective (UPLXAPO60XO, Olympus, Tokyo, Japan). Colocalization (Pearson’s correlation coefficient) of TSC2 with LAMP2 was calculated using ImageJ Coloc 2 Plugin using a single z-plane image. After removal of nuclear signals by using the image of Hoechst33342 staining, 3-5 cells were manually selected at a time and calculated.

### Live-Cell Imaging

Live-cell imaging analyses were similarly done as described previously (31). Briefly, HEK293 or HeLa cells that were seeded in a glass-bottom dish (Asahi Glass, Tokyo, Japan) were transiently transfected with CMV-R-GECO1. After 24 h, the cells were treated as follows. For carbachol stimulation, the medium was replaced with serum-free DMEM for 3 h. Time-lapse images were acquired under an FV3000 confocal laser-scanning microscope (UPLXAPO60XO, Olympus, Tokyo, Japan) before and after the addition of carbachol at 37°C. For Ionomycin stimulation, the medium was replaced with HBSS for 15 min. Time-lapse images were acquired under an FV3000 confocal laser-scanning microscope equipped with a numerical aperture oil-immersion objective (UPLXAPO40XO, Olympus, Tokyo, Japan) before and after the addition of ionomycin at 37°C.

### Statistical Analysis

Statistical analysis was performed by one-way analysis of variance (ANOVA) followed by Tukey’s test using Origin 8.0 (Micro Software, Northampton, MA, USA) or two-tailed unpaired Student’s t-test using Microsoft Exel, and p-values less than 0.05 are considered statistically significant.

## Data availability

All data described are contained within this article or the supporting information.

## Acknowledgments

We thank all members of the laboratory of Molecular and Cellular Regulation for their valuable support and discussion.

