## Supplemental Figure S1 for "Calmodulin enhances mTORC1 signaling by preventing TSC2-Rheb binding"

List of material included:

Figure S1…………page S-2

As separate files:
Movie S1
Movie S2
Movie S3
Movie S4

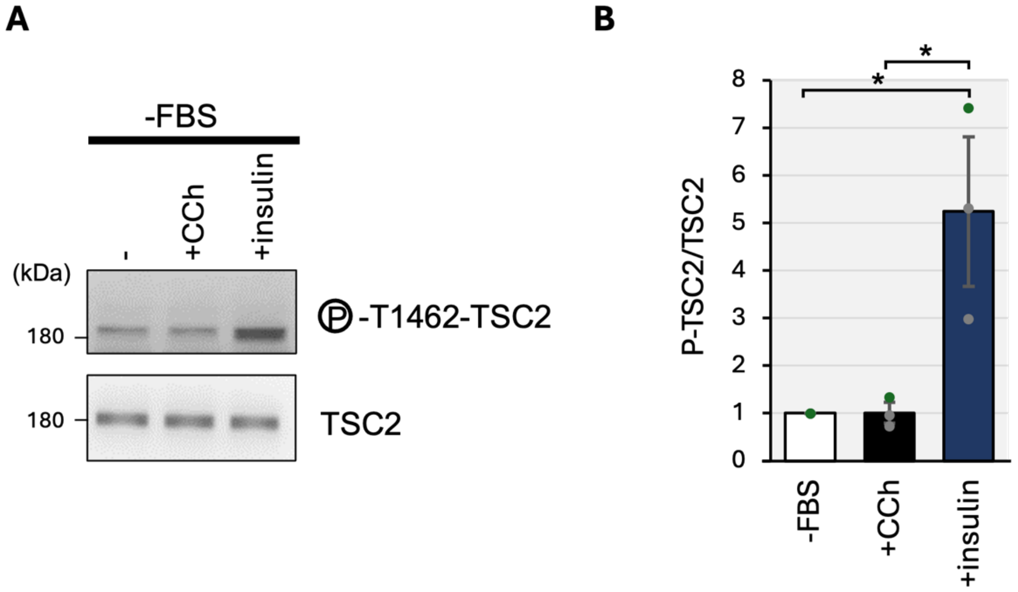


**Figure S1. Carbachol does not affect Akt-mediated phosphorylation of TSC2 at Thr1462.**

(A) HEK293 cells were starved of serum (−FBS) for 3 hr, and then stimulated with carbachol (+CCh,100 μM) or insulin (100 nM) for 15 min. Cell lysates were analyzed by Western blotting with the indicated antibodies.

(B) Quantitation of the relative intensity of phospho-Thr1462-TSC2 to total TSC2 of (A), in the FBS (-) was set to 1. Graphs represent mean ± SEM of three independent experiments, One-way ANOVA with Tukey's test, * $p < 0.05$.
